# Soil bacterial neutral lipid fatty acids: Markers for carbon storage or necromass?

**DOI:** 10.1101/2024.12.02.626346

**Authors:** Stefan Gorka, Alberto Canarini, Hannes Schmidt, Christina Kaiser

**Affiliations:** Centre for Microbiology and Environmental Systems Science, University of Vienna, Austria; University of Vienna, Doctoral School in Microbiology and Environmental Science, Vienna, Austria; Department of Biological, Geological and Environmental Sciences, University of Bologna, Italy

**Author notes:** Corresponding authors: Stefan Gorka,; Christina Kaiser.

**Keywords:** NLFA, triacylglycerol, diacylglycerol, phospholipid turnover, bacterial carbon storage, WS/DGAT

## Abstract

Carbon storage is a common strategy of soil microbes to cope with resource fluctuations. Fungi use neutral lipids (triacylglycerols, TAGs) for storage, which can be quantified via their derived fatty acids (NLFAs). NLFAs specific to bacteria can also be abundant in soils, but are rarely analysed as soil bacteria are assumed to not store TAGs. Instead, bacterial NLFAs are thought to derive from degraded phospholipids (diacylglycerols, DAGs), and thus indicate bacterial necromass, but this interpretation lacks evidence. In this perspective, we synthesise knowledge from the literature and our own experimental results on the origin of soil bacterial NLFAs. In sum, we provide evidence that bacterial NLFAs are predominantly derived from TAGs used for carbon storage: (1) Several pure culture studies provide evidence for TAG production in selected bacterial isolates. (2) Screening of genomes showed that wax ester synthase/diacylglycerol acyltransferases, which mediate the last step of TAG synthesis, are abundant in bacterial isolates from soil, suggesting a widespread genetic capability to produce TAGs. (3) We experimentally created conditions of excess labile carbon by adding isotopically labelled glucose to soil. Glucose-^13^C was rapidly allocated into bacterial NLFAs, with higher relative enrichment than phospholipid-derived fatty acids, indicating storage. (4) DAGs are not necessarily produced—and may only be intermediate compounds—during phospholipid degradation. We conclude that soil bacterial NLFAs are mainly derived from storage compounds, but a potential contribution from degraded phospholipids needs further validation. Isotopic labelling could resolve this, making NLFAs a valuable biomarker for microbial storage compounds in soil.

**Highlights:**

- Bacterial NLFAs originate from triacylglycerols (TAGs) or degraded phospholipids
- Neutral lipids are not necessarily produced during phospholipid degradation
- Soil bacteria have the genetic potential to produce TAGs for storage
- Rapid transfer of excess glucose-^13^C into soil bacterial NLFAs suggests storage
- Bacterial NLFAs are markers for carbon storage rather than necromass

## 1. Introduction

Soil organic matter represents the largest pool of terrestrial carbon, and its turnover is primarily mediated by soil microbes. Biological marker compounds provide valuable insights into the processes and pools involved in soil organic matter cycling. For instance, profiling of lipid-derived fatty acid methyl esters (FAMEs) is an established method to analyse soil microbial communities. In the FAME extraction method, lipids are usually fractionated based on their polarity. Polar phospholipid fatty acids (PLFAs) are widely used as sensitive indicators of microbial biomass and community composition (Frostegård et al., 2011; Willers et al., 2015). Additionally, the FAME extraction method can yield neutral lipid fatty acids (NLFAs), but their collection is usually restricted to fungi-specific fatty acids. Fungi produce neutral triacylglycerols (TAGs), which are composed of three fatty acids attached to a glycerol backbone. They serve as common carbon storage compounds in eukaryotic organisms and are collected as NLFAs in the FAME extraction method (Murphy, 2012; Gorka et al. 2023a). Thus, specific NLFAs associated with fungi can be used to estimate fungal carbon storage (Bååth, 2003).

Bacteria have also been shown to produce TAGs for carbon storage in culture studies (Alvarez and Steinbüchel, 2002; Wältermann and Steinbüchel, 2005), and NLFAs specific to bacteria can also be abundant in soil extracts. However, the origin of bacterial NLFAs has been interpreted in two different ways. Bacterial NLFAs have been used as markers for carbon storage, analogous to fungal NLFA (Ngosong et al., 2020; Mason-Jones et al., 2023). In contrast, some authors assume that the synthesis of TAGs in bacteria is either non-existent or not widespread, and instead suggest that bacterial NLFAs are derived from degraded phospholipids (Bååth, 2003; Rinnan et al., 2005; Amir et al., 2008; Rinnan and Bååth, 2009; Koyama et al., 2018; Honvault et al., 2021). This way, bacterial NLFAs originate from dead cells and are indicative of bacterial necromass. This interpretation not only comes with certain assumptions on phospholipid degradation pathways and extracellular lipid stabilisation, but also suggests that a certain portion of fungal NLFAs may be derived from dead fungal cells.

Both carbon storage and necromass biomarkers recently received attention as important soil carbon pools (Liang et al., 2017; Kästner et al., 2021; Mason-Jones et al., 2021). Bacterial NLFAs may serve as a very powerful biomarker, but consensus is missing whether they are reliable indicators of carbon storage or necromass. Soil microbes invest in carbon storage compounds particularly in situations of excess availability, and use them as important resources to overcome limiting conditions (Mason-Jones et al., 2021). Their analysis thus provides information on the physiological state of the microbial community. Microbial necromass has repeatedly been identified to form a large portion of soil organic carbon (Liang et al., 2019), and is usually measured as amino sugars derived from hydrolysed chitin and peptidoglycan (Joergensen, 2018; Salas et al., 2023; Salas et al, 2024).

Here, we aim to synthesise knowledge on the potential usage of soil bacterial NLFAs. We weigh the arguments for their application as carbon storage or necromass markers, discuss both interpretations critically and contrast them with our own empirical observations. We provide suggestions for future research to elucidate ambiguities in this matter, and argue that bacterial NLFAs have the potential to be an easily extractable yet powerful metric in soil science.

## 2. A short history of soil NLFA extractions and usage as fungal biomarkers

NLFAs are usually extracted concurrently with PLFAs in the FAME extraction method. It consists of total lipid extraction, and subsequent fractionation where lipids are separated based on their polarity. Lipid fractionation is generally achieved by successively eluting with chloroform (neutral lipids), acetone (glycolipids), and methanol (phospholipids) on silicic acid columns. The final step in the protocol is the derivatisation of the fractionated lipids into fatty acids (e.g. via mild alkaline methanolysis). The resulting fatty acid methyl esters are usually measured via gas chromatography. This commonly used method is based on the lipid extraction method by Bligh and Dyer (1959), adapted for microbial profiling by White et al. (1979). It was modified for soil extractions by the pH-adjustment of the monophasic chloroform:methanol:water extractant by replacement of water with citrate buffer of pH=4 (Frostegård et al., 1991), and later for high-throughput sample analysis by reducing the extractant volume and using 96-well silica column plates (Byer and Sasser, 2012). Recently it was found that the polarity of chloroform—which may be stabilised with small amounts of a more polar solvent like ethanol—is critical for the recovery of neutral lipids from silicic acid columns (Drijber and Jeske, 2019). Adding 2% ethanol to chloroform allows the complete recovery of TAGs in the neutral lipid fraction, but pure chloroform can lead to an incomplete fractionation (Gorka et al., 2023a). It is therefore important to note that previous studies which measured NLFAs in soil may have underestimated NLFA yields.

Soil NLFAs are most commonly extracted to quantify arbuscular mycorrhizal biomass via the specific NLFA 16:1ω5 (Olsson et al. 1995; Lekberg et al., 2022). The use of NLFA biomarkers of other fungal groups (ascomycete- and basidiomycete-specific fatty acids) was popularised in the conclusive paper by Bååth (2003). In this study, a labile carbon source (glucose) was added to soils, which created conditions for storage compound production as other nutrients became limiting for fungal growth. In consequence, the fungi-specific NLFAs 18:2ω6,9 and 18:1ω9 increased 60-fold and 10-fold, respectively, while no increase was observed when glucose was added together with nitrogen and phosphorus. This strongly indicates that fungal NLFAs are derived from compounds used for carbon storage. Fungi-specific NLFAs can therefore be used as indicators of fungal carbon storage, and the corresponding NLFA/PLFA ratios infer the physiological state of fungi (Rinnan et al., 2005; Rinnan and Bååth, 2009).

## 3. The enigmatic origin of soil bacterial neutral lipids

Full fatty acid profiles that include bacteria- as well as fungi-specific NLFAs are rarely considered in soil studies, even though the taxonomic specificities of marker fatty acids correspond across lipid fractions in the FAME method (Gorka et al., 2023a). We searched studies which analysed complete PLFA and NLFA soil profiles and found only 15 publications from 2002 to 2023. Closely inspecting these 15 publications showed that full NLFA profiles show significant amounts of fatty acids specific to bacteria (Table 1). We also extracted PLFA and NLFA profiles from three different soils, and found consistently high amounts of bacterial NLFAs (60-77% of nmol PLFAs g^−1^ dm) in a beech forest, spruce forest, and grassland soil (Fig. 1). In our extraction, the majority of fatty acids derived from phospholipids were also recovered in NLFA profiles, showing similar total NLFA counts compared to PLFAs (NLFA/PLFA count ratios of 0.95–1.05). Counts of bacteria-specific fatty acids were also similar between NLFAs and PLFAs (NLFA/PLFA count ratios of 0.91–1.08; Table 1). Interestingly, while almost all fatty acids specific to gram-positive bacteria were found in both PLFA and NLFA profiles, some fatty acids specific to gram-negative bacteria were only present as PLFAs (16:1ω9, 18:1ω5) or as NLFAs (17:1ω8, 19:0cy(ω9); Table S1). The values we found are in the range of previously published soil fatty acid profiles, where bacterial NLFA abundances account for 9–82% of PLFA abundances (mean 39%), and 0.15–1.14 as NLFA/PLFA count ratios (mean 0.69; Table 1). As previously mentioned, recent studies have shown that classic chloroform extraction might lead to NLFA losses, therefore these numbers might represent an underestimation.

**Figure 1.**
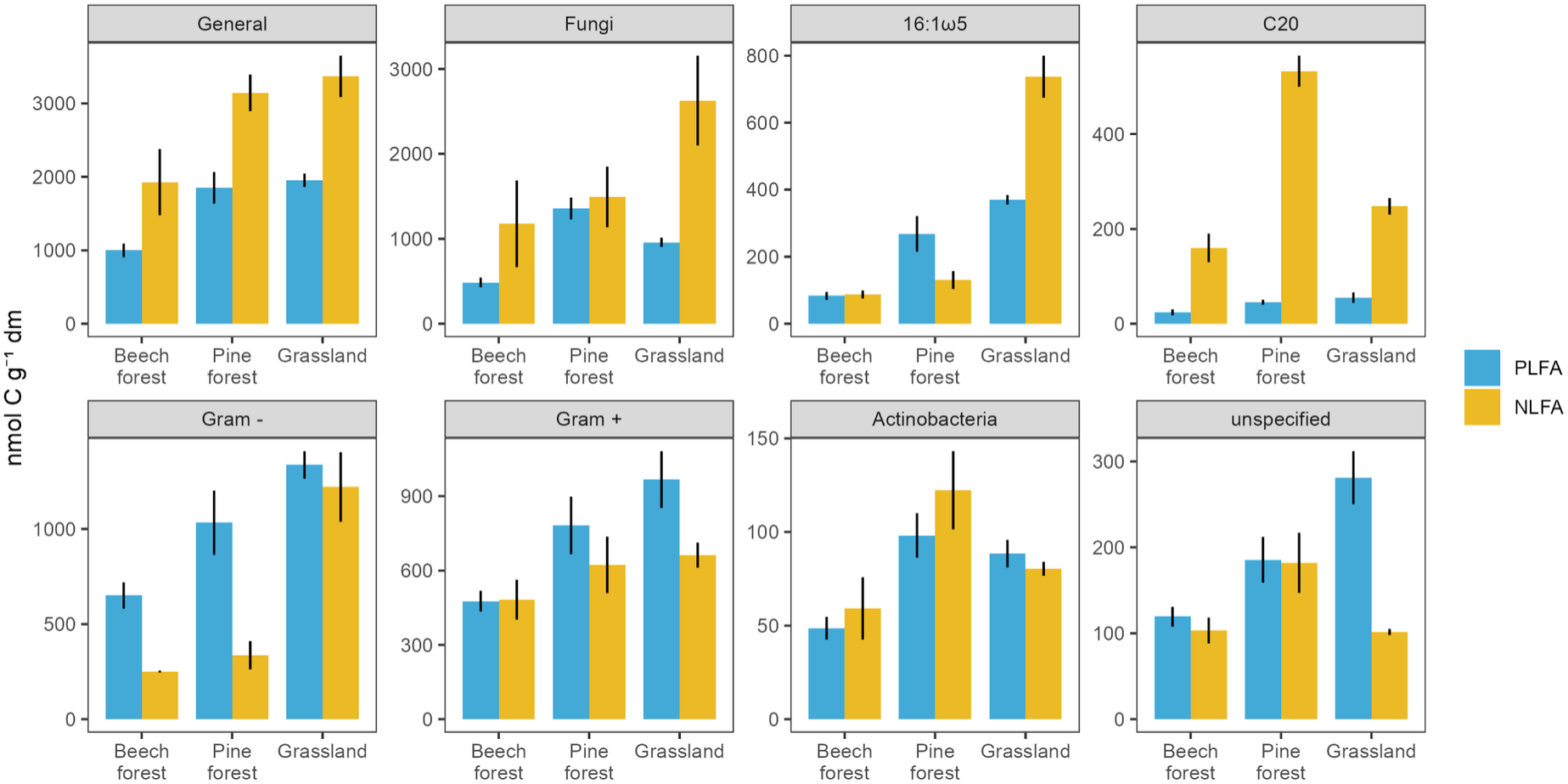
Abundances of PLFAs and NLFAs in three different soils. In all analysed soils, considerable amounts of bacteria-specific NLFAs were found. The fatty acid 16:1ω5 is depicted separately as it is specific to both arbuscular-mycorrhizal fungi and gram-negative bacteria. C20 denotes fatty acids with 20 carbon atoms. Data is presented as the sum of all fatty acids specific to each respective microbial group. Error bars represent the standard error. For a description of the sampling and extraction procedures, see *Materials and Methods* section 8.2.

**Table 1.** Comparison of full PLFA and NLFA profiles extracted from soils or similar ecological matrices. Fatty acid (FA) counts refer to the number of observed or analysed fatty acids. Abundance data were always calculated relative to mass (e.g. nmol FAs g^−1^ soil). Asterisks indicate that authors noted the given usage of bacterial NLFAs with caution, and considered the alternative interpretation as well. See *Materials and Methods* section 8.1 for details.

| Reference | Soil | Counts of bacteria-specific FAs |  |  | NLFA/PLFA abundance ratios |  |  |  |  | Usage of bacterial NLFAs |
| --- | --- | --- | --- | --- | --- | --- | --- | --- | --- | --- |
|  |  | PLFA | NLFA | NLFA/PLFA | Total | Fungi | Bacteria | Gram+ | Gram- |  |
| Hedlund et al. (2002) | Agricultural field | 13 | 2 | 0.15 | 0.72 | - | 0.33 | - | 0.66 | - |
|  | Beech forest | 13 | 2 | 0.15 | 0.27 | - | 0.09 | - | 0.17 |  |
| Bååth (2003) | Agricultural | 14 | 14 | 1 | 0.58 | 1.95 | 0.27 | 0.34 | 0.22 | Necromass |
|  | Beech forest (A-horizon) | 14 | 14 | 1 | 0.4 | 0.54 | 0.2 | 0.25 | 0.12 |  |
|  | Forest humus | 11 | 11 | 1 | 0.42 | 0.46 | 0.27 | 0.32 | 0.14 |  |
|  | Grassland | 13 | 13 | 1 | 0.48 | 0.38 | 0.11 | 0.16 | 0.09 |  |
| Malosso et al. (2004) | Antarctic soil | - | - | - | - | - | - | - | - | Necromass* |
| Rinnan et al. (2005) | Arctic heathland | 13 | 13 | 1 | 0 | 2.15 | 0.31 | - | - | Necromass |
| Bradley et al. (2007) | Grassland (sandy loam) | 7 | 1 | 0.14 | 0.31 | - | - | - | - | Necromass* |
|  | Grassland (silt loam) | 9 | 4 | 0.44 | 0.67 | - | - | - | - |  |
|  | Grassland (clay loam) | 9 | 4 | 0.44 | 0.77 | - | - | - | - |  |
| Rinnan and Bååth (2009) | Tundra heath | 18 | 10 | 0.56 | 0.61 | 1.31 | 0.26 | 0.42 | 0.14 | Necromass |
| Rinnan et al. (2013) | Tundra heath | 17 | 8 | 0.47 | 0.68 | - | - | - | - | - |
| Ferrari et al. (2015) | Agricultural soil (Summer) | - | - | - | 9.79 | - | - | - | - | - |
|  | Agricultural soil (Winter) | - | - | - | 5.25 | - | - | - | - |  |
| Koyama et al. (2018) | Nevada desert (covered vegetation) | 10 | 8 | 0.8 | 0.36 | 0.55 | 0.14 | - | - | Necromass |
|  | Nevada desert (plant interspace) | 10 | 8 | 0.8 | 0.48 | 0.72 | 0.14 | - | - |  |
| Grossman et al. (2019) | Temperate forest (experimental) | - | - | - | 1.15 | 2.22 | 0.84 | - | - | - |
| Ngosong et al. (2020) | Temperate forest | 14 | 16 | 1.14 | 0.2 | - | - | - | - | Storage |
| Honvault et al. (2021) | Rhizosheath soils | 8 | 8 | 1 | 0.98 | 0.98 | 0.67 | 0.34 | 1.02 | Necromass |
| Butler et al. (2023) | Soils from West head, Australia | - | - | - | 2.26 | - | - | - | - | Storage* |
| Gorka et al. (2023a) | Agricultural soil | 14 | 5 | 0.36 | 4.01 | 1.93 | 0.31 | 0.22 | 0.43 | Storage* |
|  | Beech forest | 11 | 12 | 1.09 | 1.05 | 1.32 | 0.71 | 1.05 | 0.47 |  |
|  | Grassland | 15 | 5 | 0.33 | 3.68 | 2.31 | 0.81 | 0.52 | 1.1 |  |
| Gorka et al. (2023b) | Beech forest | 13 | 13 | 1 | 0.97 | 0.76 | 0.82 | 1.29 | 0.53 | Storage* |
|  | **Mean** | **12.3** | **8.55** | **0.69** | **1.5** | **1.26** | **0.39** | **0.49** | **0.42** |  |
| *This publication* | Beech forest | 12 | 13 | 1.08 | 1.33 | 2.42 | 0.69 | 0.39 | 1.01 |  |
|  | Pine forest | 11 | 10 | 0.91 | 1.14 | 1.1 | 0.6 | 0.33 | 0.8 |  |
|  | Grassland | 11 | 10 | 0.91 | 1.4 | 2.74 | 0.77 | 0.91 | 0.69 |  |
|  | **Mean** | **11.3** | **11** | **0.96** | **1.29** | **2.09** | **0.69** | **0.54** | **0.83** |  |

Where do these bacterial NLFAs come from? In the literature, they have been interpreted in two different ways: (i) bacterial NLFAs indicate storage compounds, and/or (ii) they indicate bacterial necromass (Fig. 2). However, the majority of studies which measured bacterial NLFAs in soils used them as markers for necromass (Table 1). The underlying reasoning is that bacterial cells (presumably) do not contain TAGs, and when a cell dies, phospholipids are released and undergo rapid degradation (Bååth, 2003; Zhang et al., 2019). The initial step of this process involves the cleavage of the charged phosphate head, leading to the formation of a neutral diacylglycerol (DAG), which may then be stabilised in soil for an unknown amount of time. Using bacterial NLFAs as necromass biomarkers implicitly assumes that DAGs will—similar to TAGs—elute in the neutral lipid fraction of the FAME protocol, even though they are slightly more polar than TAGs due to the additional hydroxyl group. This assumption has never been verified to the best of our knowledge. Consequently, we tested the recovery of fatty acids derived from a DAG in the FAME extraction method. Our results show that both, TAG and DAG, co-elute in the neutral lipid fraction (Table 2; Gorka et al., 2023a), which confirms that potential remains of former phospholipids in the soil will be measured as NLFAs in the FAME extraction method. Whether they are representative of bacterial necromass, however, also depends on how stable phospholipid-derived DAGs are in soils, a point which we discuss in section 4.2. In contrast, use of bacterial NLFAs as markers for carbon storage is supported by multiple observations of bacterial TAG storage in intracellular lipid droplets (Wältermann et al., 2005; Wältermann and Steinbüchel, 2005; Wältermann and Steinbüchel, 2006). These are usually limited to bacterial isolates, and are predominantly found in Actinobacteria (Barksdale and Kim, 1977; Arabolaza et al., 2008; Hernández and Alvarez, 2010; Cortes and de Carvalho, 2015; Eberly et al., 2013; Röttig et al., 2016). However, the prevalence of bacterial TAG production in soils is unknown. We provide a more detailed exploration of this in section 5.1.

**Figure 2.**
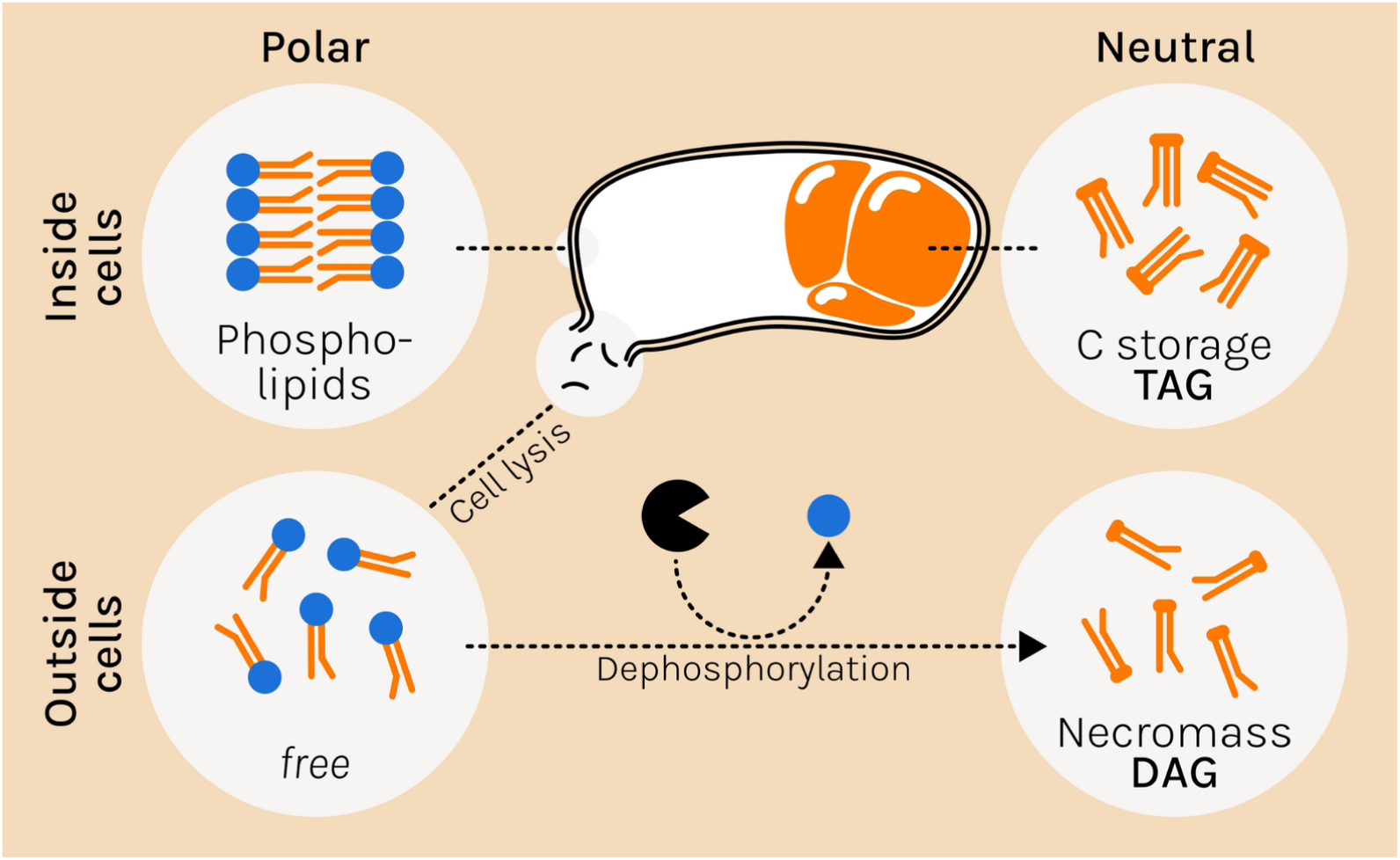
Soil bacterial NLFAs have been interpreted in two ways: (i) They are derived from intracellular TAGs and are indicative of carbon storage. (ii) They are derived from membrane phospholipids, which are decomposed to DAGs by extracellular phosphatases after cell death, and thus indicative of bacterial necromass. Both TAGs and DAGs would be collected in the neutral lipid fraction in the FAME method, and are measured as NLFAs.

**Table 2.** Recovery of fatty acids derived from a diacylglycerol (DAG), and the fatty acid and alcohol moieties of a wax ester (WE) in the three lipid fractions of neutral lipids (NL), glycolipids (GL) and phospholipids (PL). Pure lipid standards were fractionated on silica gel columns and derivatised via mild alkaline methanolysis. The recovery was calculated by comparison to unfractionated samples, and values are presented in % recovery. DAG, n=3; WE, n=4; n.d., not detected. See *Materials and Methods* section 8.4 for a detailed methodology.

| Lipid fraction | % Recovery |  |  |
| --- | --- | --- | --- |
|  | DAG-fatty acids | WE-fatty acid | WE-alcohol |
| NL | 99.3 | 78.9 | 69.6 |
| GL | 0.7 | 1.4 | n.d. |
| PL | 0.2 | 0.3 | n.d. |

Both interpretations of bacteria-specific NLFAs are theoretically plausible and not mutually exclusive. This problem is exemplified when comparing microbial pure culture and soil fatty acid profiles. In a previous publication, we extracted PLFA and NLFA profiles from pure cultures of bacteria and fungi (Gorka et al., 2023a). These profiles were primarily separated by taxa, regardless of whether the fatty acids came from phospholipids or neutral lipids (PERMANOVA P < 0.001, R^2^ = 0.52; Fig. 3a, Table 3a). PLFAs and NLFAs were also separated, but with a much lower explanatory power (P < 0.001, R^2^ = 0.07).. This not only shows that bacteria produce neutral lipids—likely TAGs—but also that microbes use the same fatty acids for phospholipids and neutral lipids and in similar proportional distribution in the two fractions. However, in the soil environment this interpretation is more complicated. In contrast to the pure culture profiles, the PLFA and NLFA profiles of soils were primarily separated by the lipid type (Fig. 3b), and PERMANOVA analysis showed similar contributions of the lipid source (P < 0.001, R^2^ = 0.33) and soil type (P < 0.001, R^2^ = 0.30; Table 3b). At a first glance this data could be interpreted as indicative of NLFA representing necromass rather than carbon storage since the soil fatty acid profiles do not reflect the pure culture profiles. However, from the comparison of soil PLFA and NLFA profiles one cannot conclusively infer the origin of NLFAs. If bacteria produce TAGs, we would expect different relative concentrations of NLFAs to PLFAs, as storage production depends on the overall physiological state of the microbial community, which is linked to environmental nutrient limitations (Bååth et al., 2003; Wältermann et al., 2005). Moreover, there may be a preferential allocation of specific fatty acids into storage lipids. For example, this is the case in arbuscular mycorrhizal fungi which preferentially allocate the fatty acid 16:1ω5 into TAG (Olsson et al., 1995). On the other hand, if bacterial neutral lipids are derived from decomposed phospholipids (i.e. DAGs), the NLFA concentration would depend on the rate of phosphorylation, microbial turnover, and the retention of DAG, potentially leading to different concentrations of PLFAs and NLFAs. Thus, neither interpretation can be dismissed nor proved by only examining the relative distributions of the fatty acids.

**Figure 3.**
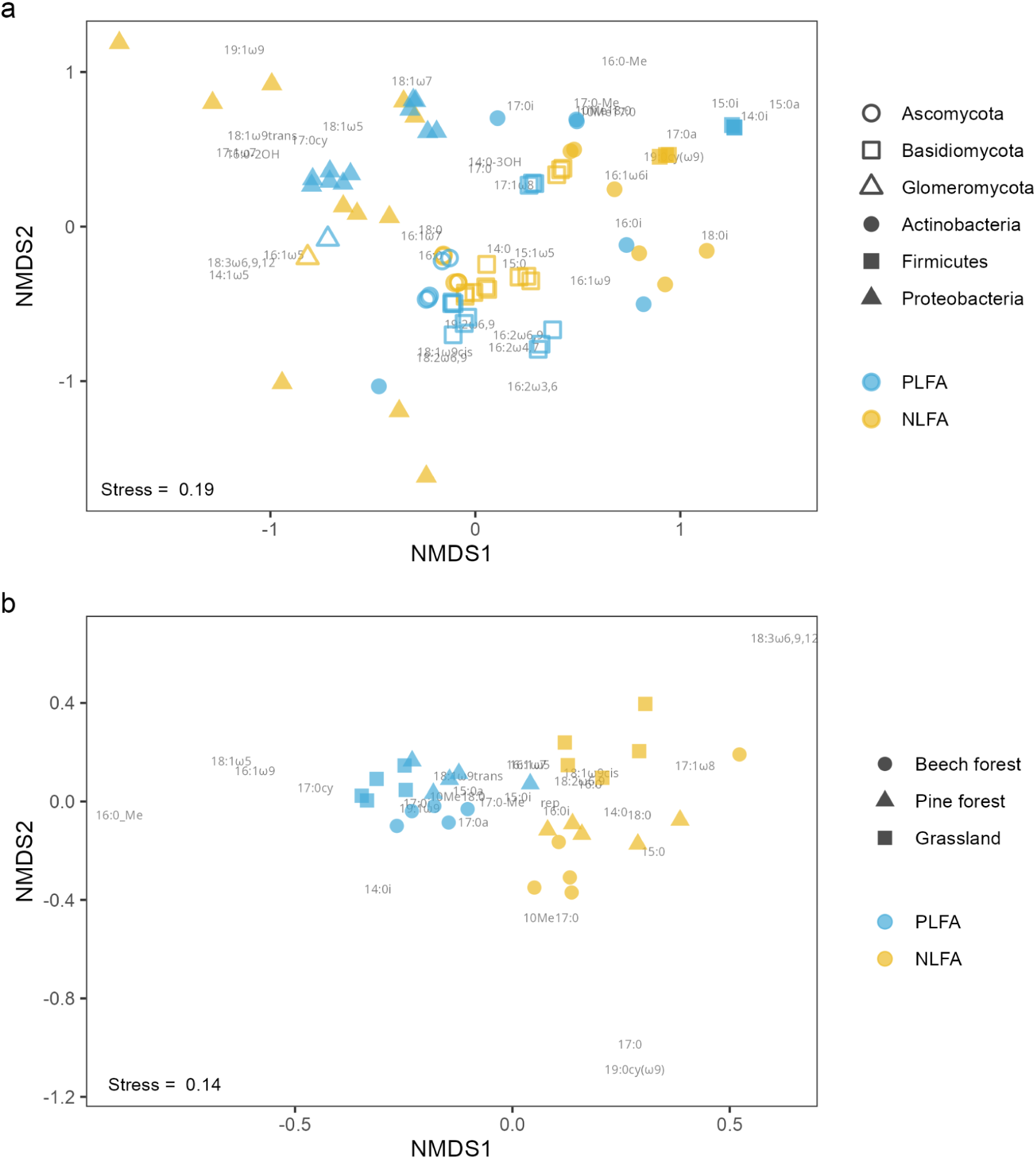
Non-metric multidimensional scaling (NMDS) plots of PLFA and NLFA profiles from **(a)** pure culture strains (data from Gorka et al., 2023a), and **(b)** three different soils (same data as Fig. 1). While pure culture profiles primarily cluster by phylum, soil profiles primarily cluster by lipid type (Table 3). Data provided in (a) mol%, and (b) nmol C g^−1^ dm. Individual points represent replicate samples. For a description of the sampling and extraction procedures, see *Materials and Methods* section 8.2.

**Table 3.** Permutational multivariate analysis of variance (PERMANOVA) results of fatty acid profiles from **(a)** pure culture strains, and **(b)** soil profiles (see Fig. 3). ‘Lipid type’ refers to PLFA vs. NLFA. df, degrees of freedom; SS, sum of squares.

| **a** | df | SS | R<sup>2</sup> | F | P |
| --- | --- | --- | --- | --- | --- |
| **Lipid type** | 1 | 1.607 | 0.075 | 12.785 | 0.001 |
| **Taxon** | 5 | 11.223 | 0.522 | 17.863 | 0.001 |
| **Residual** | 69 | 8.67 | 0.403 |  |  |
| **Total** | 75 | 21.50 | 1 |  |  |

| **b** | df | SS | R<sup>2</sup> | F | P |
| --- | --- | --- | --- | --- | --- |
| **Lipid type** | 1 | 0.838 | 0.330 | 23.165 | 0.001 |
| **Soil** | 2 | 0.759 | 0.299 | 10.483 | 0.001 |
| **Residual** | 26 | 0.941 | 0.371 |  |  |
| **Total** | 29 | 2.538 | 1 |  |  |

## 4. Bacterial NLFAs as necromass markers

Soil microbial necromass is an important contributor to soil organic matter, accounting for approximately 50% of total soil organic carbon (Liang et al., 2017; Liang et al., 2019; Wang et al. 2021). Amino sugars, which are found in cell walls as chitin and peptidoglycan, are usually used to quantify microbial necromass (Kästner et al., 2021; Salas et al., 2024). They allow estimates of fungal and bacterial necromass, but deeper taxonomic analysis is restricted as there are only four types of necromass-specific amino sugars, with only muramic acid being unique to bacterial peptidoglycan (Joergensen, 2018).

Apart from chemically recalcitrant cell wall fragments, dead cells leach a diverse set of cytoplasmic and membrane molecules (Camenzind et al., 2023). These are continually decomposed from polymers to monomers, and their stabilisation is intricately linked to the spatial heterogeneity and temporal variability of the soil matrix, which determine their accessibility to soil microbes (Lehmann and Kleber, 2015; Lehmann et al., 2020). Furthermore, the chemical nature of the decomposition products (e.g. polarity, ionisation) is itself a factor for stabilisation (von Lützow et al., 2006). Lipids may therefore form a significant part of necromass that is usually not measured.

Soil bacterial NLFAs may give valuable insight into membrane-lipid derived bacterial necromass at a certain stage of its decomposition. However, in order for NLFAs to be derived from decomposed phospholipids (DAGs), it is a prerequisite that (i) phospholipids derived from dead cells are decomposed extracellularly into DAGs, and (ii) these DAGs are stabilised in a recalcitrant manner (physico-chemically or spatially). In the following, we discuss current knowledge on these aspects.

### 4.1 Decomposition pathways of soil phospholipids

Soil phospholipids are rapidly degraded within hours to days (White et al., 1979; Janzen et al., 1994; Zhang et al., 2019). The interpretation that soil NLFAs are derived from degraded phospholipids (DAGs) assumes that the majority of phospholipids is hydrolysed by a particular class of phosphodiesterases. These enzymes belong to the phospholipase C family, which cleave between the glycerol and the phosphate group (Titball, 1993). Other proteins in the phospholipase superfamily cleave phospholipid molecules at various positions, which leads to the formation of distinct breakdown products other than DAGs (Fig. 4a; Fisher and Jain, 2009; Ramrakhiani and Chand, 2011; Barman et al., 2018). Phospholipases A_1_ and A_2_ hydrolyse the fatty acyl-ester bonds at the respective *sn*-1 and *sn*-2 positions of the glycerol moiety, producing free fatty acids and lysophospholipids (Köhler et al., 2006, Sitkiewicz et al., 2007). Phospholipase B cleaves both fatty acids, producing free fatty acids and the phosphorylated head group attached to glycerol-3-phosphate. Phospholipase D cleaves after the phosphate, producing a polar phosphatidic acid and an unphosphorylated head group. Continuing this pathway, phosphatidic acid can further be dephosphorylated to generate DAG (Pyne et al., 2004; Carman and Han, 2006; Fisher and Jain, 2009). Whether extracellular DAGs are produced is therefore determined by the relative abundances of the various phospholipases in soil, where only phospholipase C, as well as downstream phosphorylation by phospholipase D, can yield DAGs from phospholipids (Fig. 4b).

**Figure 4.**
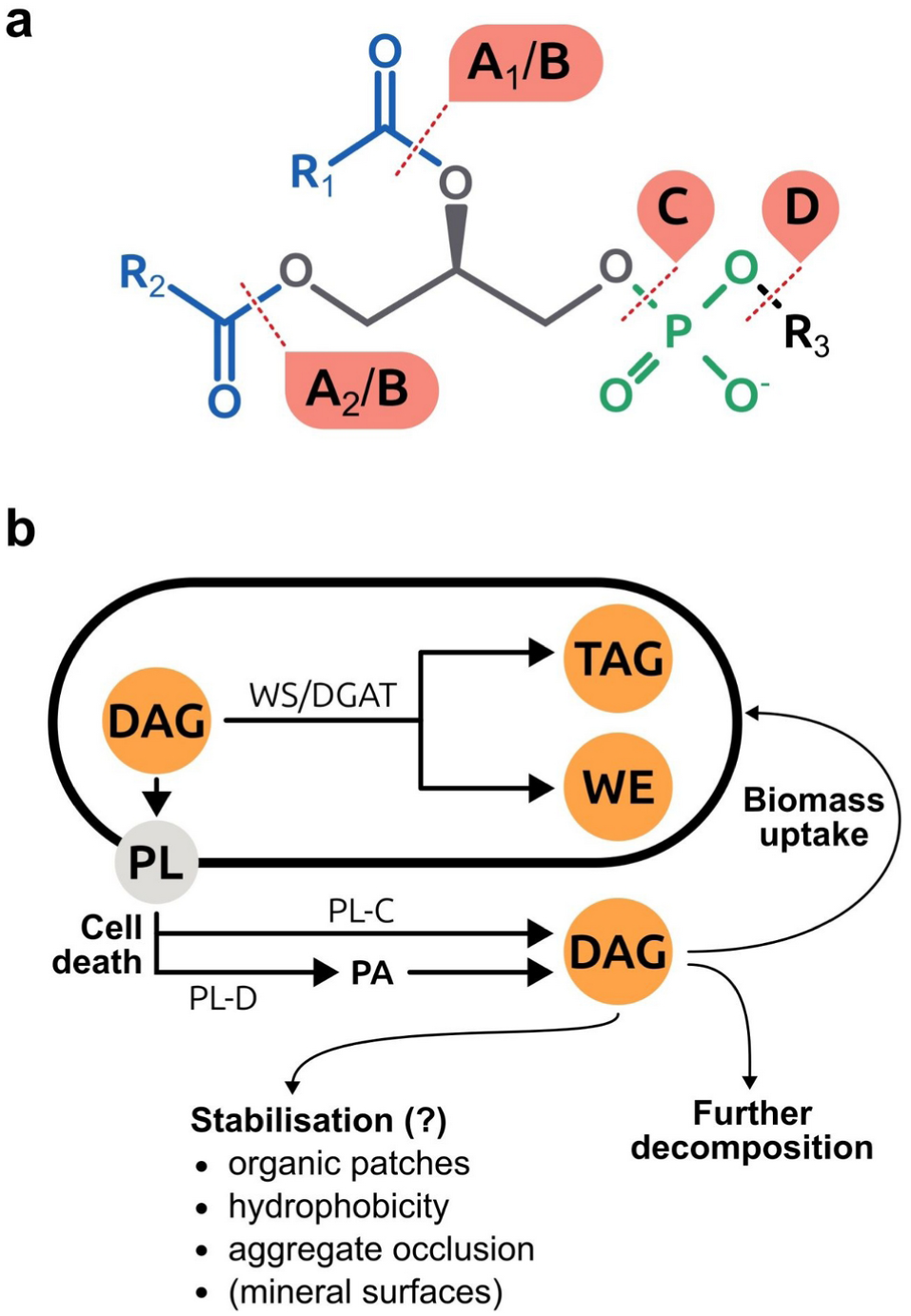
Potential sources of bacterial neutral lipids in soils. Pathways for degradation of phospholipids that lead to DAG formation. **(a)** Possible sites of hydrolytic cleavage by phospholipases on a phospholipid. The letters in the red boxes show the respective phospholipase families. **(b)** Synthesis and decomposition pathways that lead to intracellular and extracellular neutral lipids, which are possible sources of soil bacterial NLFAs (marked in orange). TAG, triacylglycerols; WE, wax esters; DAG, diacylglycerols; PL, phospholipids; WS/DGAT, wax ester synthase/diacylglycerol acyltransferase; PL-C and -D, phospholipase C and D; PA, phosphatidic acid.

All phospholipase classes can be secreted by bacteria (Stinson and Hayden, 1979; Flieger et al., 2000; Hagishita et al., 2000; Farn et al., 2001; Sugiyama et al., 2002; Simkhada et al., 2007; Sugimori et al., 2012), and phosphodiesterases have been suggested to be widespread among soil bacteria (Kuroshima and Hayano, 1982; Lidbury et al., 2017). However research is lacking on their relative contributions to soil phospholipid turnover. Only in a recent study, Wasner et al. (2023) found direct uptake of phospholipid-derived, organic phosphorus in an ^33^P radio-isotope incubation experiment. Since transporters have only been identified for glycerol-3-phosphate, the authors speculate that a portion of phospholipid breakdown occurs through a pathway that produces glycerol-3-phosphate as an end product. This pathway is likely to involve phospholipase D, as well as phospholipase A or B, which suggests that phospholipids are consecutively degraded by several phospholipase classes.

DAGs can further be hydrolysed by DAG- and monoacylglycerol lipases into glycerol and free fatty acids (Rengachari et al., 2012; Yuan et al., 2016; Li et al., 2020). In a previous experiment, we found that a freeze-thawing event, which likely killed a large part of the soil microbial community, decreased both fungal and bacterial NLFAs within days (Gorka et al., 2023b). This either suggests phospholipase pathways which did not yield DAGs, or further decomposition of phospholipid-derived DAGs. Another alternative explanation is that bacterial NLFAs are derived from TAGs used for storing carbon in living biomass which degraded after cell death. Although DAGs can theoretically be produced during phospholipid degradation, they are a potentially intermediate compound in only two of several pathways during phospholipid degradation. We conclude that a more systematic investigation of soil phospholipid degradation products is needed to assess their contribution to soil bacterial NLFA pools.

### 4.2 Stability of neutral lipids in soils

It is unclear how long extracellular neutral lipids persist in soil, but there are indications that they may be stabilised. Parts of stable microbial necromass can persist as lipids (Kallenbach et al., 2016; Hall et al., 2020), possibly by interaction with mineral surfaces (Olivelli et al., 2020; Neurath et al., 2021). Indeed, lipids derived from bacterial extracellular polymeric substances (EPS) have been found to attach to goethite heterogeneously in lipid-rich spots (Liu et al., 2013). While lipids are only minor components of EPS, goethite-adsorbed EPS consisted of >25% lipids, suggesting preferential adsorption of lipids to goethite. Nonetheless, since DAGs are only slightly polar, their stabilisation via organo-mineral interactions is hypothetically minor due to weak adsorption. The -CH and -OH groups of DAGs are electrostatically neutral in typical soil pH ranges, which makes them less mobile in aqueous solutions and less likely to bond to mineral surfaces or metal cations (Kleber et al., 2015). Organic matter binds to minerals in clusters, indicating that new necromass compounds preferentially associate with existing organic matter attached to mineral sites (Vogel et al., 2014). It has thus been suggested that organo-organo interactions might be another form of necromass stabilisation (Buckeridge et al., 2020; Buckeridge et al., 2022).

The hydrophobic character of lipids plays a role in protection from mineralisation (Kästner et al., 2021). Hydrophobicity has been positively linked to carbon sequestration as it reduces surface wettability and thus accessibility of organic matter to microorganisms (Piccolo et al., 1999; von Lützow et al., 2006; Song et al., 2013). Hydrophobicity also increases aggregate stability, which stabilises organic matter spatially via occlusion or clogging of intra-aggregate pores (von Lützow et al., 2006; Hafida et al., 2007; Totsche et al., 2018). Furthermore, mineralisation of reduced organic compounds like lipids is highly limited in anaerobic conditions, which can result in their preservation (Keiluweit et al., 2017). This suggests that neutral lipids may accumulate particularly in water-inaccessible pores as organic patches, and anaerobic microsites within aggregates.

Phospholipid-derived DAGs are only measured in the NLFA fraction if sufficient amounts are stabilised in soil. While literature on DAG stabilisation is limited, some studies analysed the retention of TAGs in soil which theoretically behave similarly. Soliman and Radwan (1981) recovered 62% of a TAG months after its addition to soils. However, the measured TAG may have been extracted from microbial biomass as the authors could not differentiate between added and newly produced TAGs. In a recent study, mineralisation of ^13^C-labelled plant-derived lipids started within minutes, while 36% of lipid-carbon were recovered after 15 days in grassland soil, which included significant losses of TAGs (Warren and Butler, 2023). This indicates rapid degradation of the added lipids, but does not preclude possible retention mechanisms for longer time periods. Indeed, ^14^C abundances in PLFAs were found to be close to atmospheric values, while neutral lipid extracts were depleted in ^14^C (Rethemeyer et al., 2004). This suggests that the neutral lipid pool may include older carbon, indicating either recalcitrance or slower turnover.

In sum, neutral lipids, including DAGs, are rapidly degraded, but may in part be stabilised through mechanisms such as mineral interactions, hydrophobicity, and protection in microsites. However, TAGs leached from dead cells could also persist under these conditions, which implies that the presence of NLFAs in soil could reflect both microbial carbon storage and necromass.

## 5. Bacterial NLFA as carbon storage markers

Production of carbon storage compounds is common in microorganisms (Lee, 1996; Elbein et al., 2003; Rontani, 2010; Wilson et al., 2010; Dalpé et al., 2012; Murphy, 2012; Cifuente et al., 2019; Kumar et al., 2020). They can be readily mobilised to meet energy or carbon demands under limiting conditions. Two strategies for storage can be differentiated: (i) reserve storage, where resources are continuously allocated into storage compounds at the cost of other metabolic demands, and (ii) surplus storage, where only abundant resources are allocated into storage compounds (Mason-Jones et al., 2021). While reserve storage comes with trade-offs against other physiological demands, it can be a valuable investment into future survival and reproduction. Surplus storage allows greater stoichiometric buffering and use of carbon resources, which can be particularly advantageous in environments of fluctuating resources (Manzoni et al., 2021).

Bacteria can use a wide spectrum of compounds to store carbon and energy: PHAs, glycogen, trehalose, wax esters, and TAGs are all used for carbon storage. PHAs and wax esters are unique storage compounds in bacteria and have been found in several phyla, predominantly in Proteobacteria (PHAs) and Actinobacteria (wax esters; Dalal et al., 2010; Murphy, 2012; Sehgal and Gupta, 2020). Glycogen, trehalose, and TAGs also occur in fungi and are not unique to bacteria (Preiss, 1984; Genet et al., 2000; Rangel-Castro et al., 2002; Elbein et al., 2003; Wilson et al., 2010; Mason-Jones et al., 2021). In contrast to glycogen and trehalose, TAGs are hydrophobic which allows compact storage in lipid droplets with higher energy density. Although TAGs have been observed in several bacterial species (particularly Actinobacteria, see section 5.1), their abundance in soil bacterial communities is still unclear. Below we discuss the potential across known bacterial species for TAG production and also provide experimental evidence.

### 5.1 Use of triacylglycerols (and wax esters) for carbon storage in soil bacteria

Although other forms of carbon storage like PHAs (Mason-Jones et al., 2019; Kumar et al., 2020) or glycogen (Sekar et al., 2020; Wang et al., 2020a) are assumed to occur more prevalently, TAGs have been observed in several bacterial species—particularly Actinobacteria (Alvarez and Steinbüchel, 2002). TAGs are stored in intracellular lipid droplets (Packter and Olukoshi, 1995; Alvarez et al, 1996), and their hydrophobic nature allows the accumulation of large intracellular reservoirs without affecting the osmolarity of the cytoplasm (Alvarez and Steinbüchel, 2002). Accumulation of TAGs in bacterial cultures can be stimulated if nitrogen availability is limiting growth and carbon is available in excess, and occurs predominantly during the stationary growth or sporulation phase (Hoskisson et al., 2001; Alvarez and Steinbüchel, 2002; Wältermann and Steinbüchel, 2006). Some bacteria are able to accumulate considerable amounts of TAGs when grown on carbon-rich media. For instance, *Micromonospora echinospora* was found to accumulate up to 8% TAGs of their cellular dry weight (Hoskisson et al., 2001), *Rhodococcus opacus* strain PD630 up to 76% TAGs (Wältermann et al., 2000), or 87% fatty acids (the majority of which were TAG-derived; Alvarez et al., 1996), and *Gordonia* sp. 29-96% total lipids on various growth media with significant amounts of TAGs (5-58 mg L^−1^ medium; Gouda et al., 2008).

Organisms that accumulate TAGs for carbon storage typically use specific DAG acyltransferases (DGAT) to catalyse the final addition of the third fatty acid moiety to the hydroxyl group on the DAG glycerol backbone. Contrary to eukaryotes, which use acyltransferases from two diverse DGAT families (Liu et al., 2012), bacteria preferentially utilise a specialised wax ester synthase/diacylglycerol acyltransferase (WS/DGAT, Fig. 4b; Kalscheuer and Steinbüchel, 2003; Röttig and Steinbüchel, 2013; Alvarez, 2016). As the name indicates, WS/DGAT can accept DAGs for TAG synthesis (i.e., DAG plus fatty acyl-coenzyme A, resulting in a TAG), as well as fatty alcohols to produce wax esters (i.e., fatty alcohol plus fatty acyl-coenzyme A, resulting in a wax ester).

Wax esters represent a potentially additional source of NLFAs apart from TAGs, as they consist of a fatty alcohol and a fatty acid component (Alvarez, 2016). We tested the recovery of fatty acids derived from a wax ester in the FAME extraction method. We found that wax esters co-elute in the neutral lipid fraction with relatively high recovery (ca. 80%), and the fatty acid moiety is methylated to a FAME in the derivatisation step (Table 2). Wax esters will therefore be measured as NLFAs in the FAME extraction method. We also found that the alcohol derived from a pure wax ester standard can be identified as a distinct peak in NLFA chromatograms (Fig. S1) and has reasonable recovery (ca. 70%; Table 2). Thus, it should be possible to estimate the proportion of wax esters in the total soil neutral lipid pool by comparing the total concentrations of alcohols in the neutral lipid fraction (assuming that all of them are derived from wax esters) to NLFAs. We did not, however, detect any alcohols in soil NLFA extracts (in data presented in this publication, as well as past extractions), which suggests that wax esters are not widely used for storage by soil microbes.

To estimate the potential of soil bacteria to produce TAGs (and potentially wax esters), we employed the Joint Genome Institute’s (JGI) *Integrated Microbial Genomes with Microbiome Samples Expert Review* (IMG/M-ER; Chen et al., 2023; https://img.jgi.doe.gov/) database of bacterial isolate genomes to search for protein families from the Pfam database (Paysan-Lafosse et al., 2023; https://www.ebi.ac.uk/interpro/). We specifically searched for WS/DGAT (C-terminal domain; pfam06974) and DAGAT (pfam03982) as marker enzymes for potential TAG production. The search was restricted to all bacteria that occur in the specified ecosystem type ‘soil’. Since datasets of bacterial isolates cannot fully portray actual soil bacterial communities, we additionally tested how representative the search in the JGI dataset is by estimating the occurrence of the storage compound PHA which is considered common in bacteria. For this, we searched for the PHA synthesis regulator PhaR via the PHB/PHA accumulation regulator DNA-binding domain (pfam07879; Maehara et al., 2002) as a proxy for PHA–producing bacteria. The search resulted in a total dataset of 2926 unique bacterial isolates, of which 918 were assigned to WS/DGAT, 74 to DAGAT, and 790 to PhaR.

Based on this dataset, the majority of soil Actinobacteria (66%) and a significant portion of Proteobacteria (19.2%) encode for WS/DGAT (Fig. 5, Table 4). Actinobacteria showed WS/DGAT occurrences in the class Actinomycetia (66%, n=1065), but also in Thermoleophilia (100%, n=7) and the singular species of Rubrobacteria in the dataset (Table S2). Indeed, TAG production has been observed in Actinobacteria previously in the genera *Dietzia* (Alvarez, 2003), *Gordonia* (Alvarez, 2003; Eberly et al., 2013), *Micromonospora* (Hoskisson et al., 2001), *Mycobacterium* (Akao and Kusaka, 1976; Maurya et al., 2019), *Nocardia* (Alvarez, 2003; Röttig et al., 2016), *Rhodococcus* (Alvarez, 1996; Alvarez et al., 2000; Wältermann et al., 2000, Hernández and Alvarez, 2010; Cortes and de Carvalho, 2015; Röttig et al., 2016), and *Streptomyces* (Arabolaza et al., 2008; Röttig et al., 2016). All these genera were also present in the JGI dataset and encoded for WS/DGAT.

**Figure 5.**
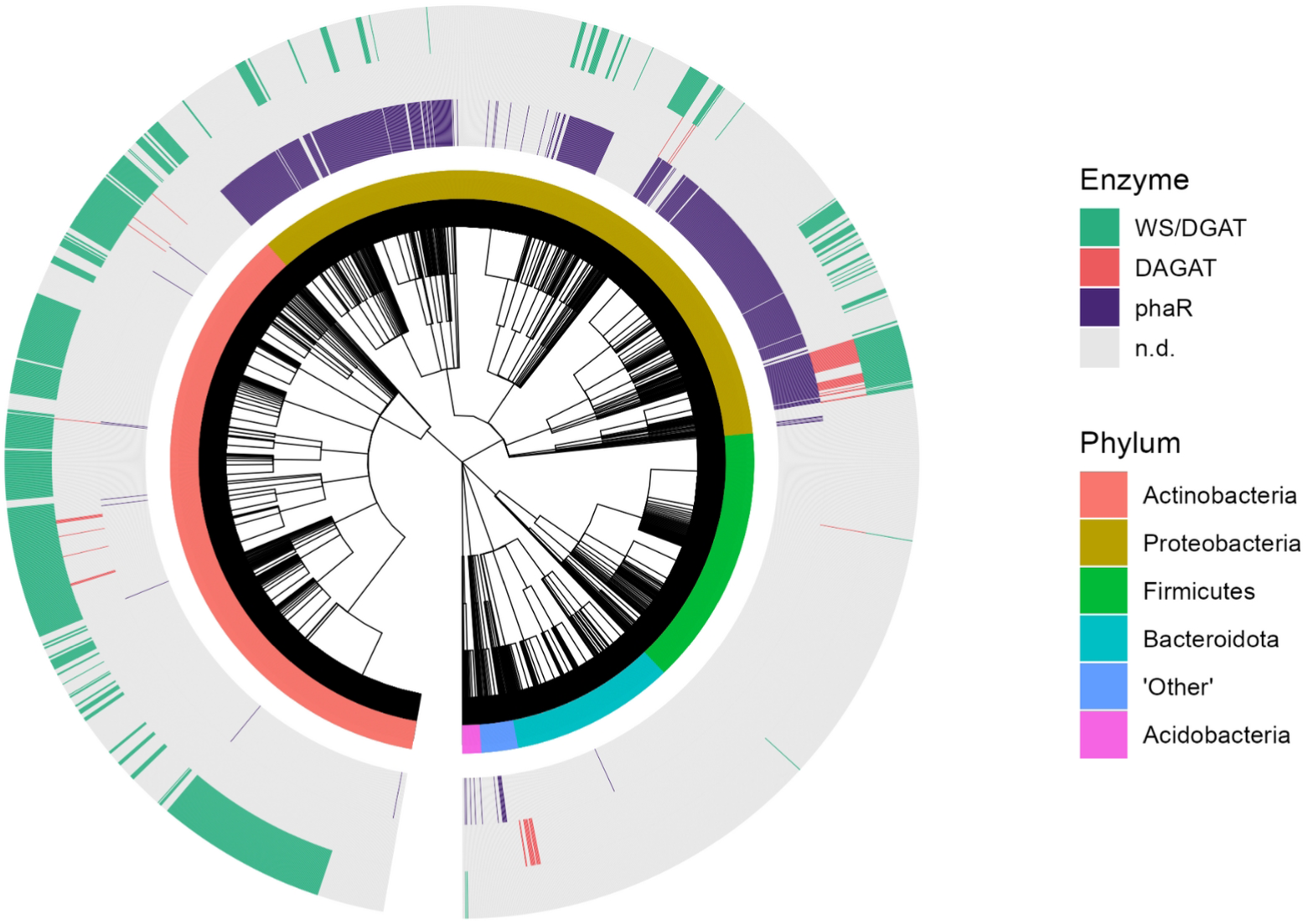
Occurrence of wax ester synthase/diacylglycerol acyltransferase (WS/DGAT), diacylglycerol acyltransferase (DAGAT), and the polyhydroxyalkanoate synthesis regulator PhaR in soil bacterial isolates based on the IMG/M-ER database. Both WS/DGAT and DAGAT catalyse the last step in TAG synthesis. ‘Other’ denotes multiple phyla with species coverage (n < 20). The cladogram shows hierarchical relationships of soil bacteria. For a detailed description of the methodology, see *Materials and Methods* section 8.5.

**Table 4.** Occurrence of specific genes for wax ester synthase/diacylglycerol acyltransferase (WS/DGAT), diacylglycerol acyltransferase (DAGAT), and the polyhydroxyalkanoate synthesis regulator PhaR in phyla of soil bacterial isolates (Fig. 5). Both WS/DGAT and DAGAT catalyse the last step in TAG synthesis. ‘Other’ denotes multiple phyla with low species coverage (n < 20).

| Phylum | n | WS/DGAT(%) | DAGAT (%) | PhaR (%) |
| --- | --- | --- | --- | --- |
| Actinobacteria (G+) | 1073 | 66.4 | 1.2 | 0.8 |
| Proteobacteria (G-) | 1054 | 19.2 | 4.5 | 72.9 |
| Firmicutes (G+) | 434 | 0.5 | 0.2 | - |
| Bacteroidota (G-) | 273 | - | - | 0.4 |
| ‘Other’ | 61 | - | 21.3 | 11.5 |
| Acidobacteria (G-) | 31 | 6.5 | - | 16.1 |

We also found substantial occurrences of WS/DGAT in Proteobacteria in the classes Deltaproteobacteria (73%, n=90), Gammaproteobacteria (16%, n=333), Betaproteobacteria (17%, n=287), and Alphaproteobacteria (10%, n=335). TAG production was observed previously in the Gammaproteobacteria *Acinetobacter* (Scott and Finnerty, 1976) and *Alcanivorax* (Kalscheuer et al., 2007); both genera also encoded WS/DGAT in the JGI dataset. Occurrences of WS/DGAT were also found in Acidobacteria in the class Acidobacteriia (7.1%, n=28), but were low to non-existent in Firmicutes (0.3% in Bacilli, n=368; 2.3% in Clostridia, n=43). Although the DAGAT family is usually associated with eukaryotes, we found occurrences in Cyanobacteria (87%, n=15) and Deltaproteobacteria (49%, n=90), and in a few species of Actinomycetia (1.2%, n=1065) and Gammaproteobacteria (1%, n=333; Table S2).

The highest occurrences of PhaR were found in Proteobacteria (73%), and fewer in Acidobacteria (16%) and phyla with low coverage in the JGI dataset (n < 20, denoted as ‘Other’; 11.5%). All other bacterial groups showed only few occurrences of PhaR (< 1%; Fig. 5, Table 4). This is consistent with available literature on PHA producing bacterial isolates (Wang and Bakken, 1998; Kadouri et al., 2005; Dalal et al., 2010; Sehgal and Gupta, 2020).

This search of soil bacterial isolate genomes certainly comes with biases (overrepresentation of culturable species, annotation reliance on known protein families), but it nonetheless provides insight on the potential use of lipids in bacterial carbon storage. We identified WS/DGAT in classes of both gram-positive bacteria (Actinobacteria) and gram-negative bacteria (Proteobacteria, Acidobacteria), and found that certain bacterial classes lacked WS/DGAT in most of their species (Firmicutes, Bacteroidota; Fig. 5, Table 4). Intriguingly, we found similar occurrences of WS/DGAT compared to phaR (31% and 27% of all analysed genomes respectively), suggesting that bacterial TAG production may be as common as PHA production. Overall, these results indicate that the potential for production of TAGs may be more widespread in soil bacteria than previously assumed.

### 5.2 Abundant labile carbon is readily incorporated into both soil PLFAs and NLFAs

We experimentally tested whether bacteria utilise neutral lipids for carbon storage by creating conditions of excess abundance of a labile carbon source. Specifically, we added ^13^C-labelled glucose to sieved beech forest soil (4 mg glucose g^−1^ soil). We then incubated the soil at a constant temperature of 20 °C, and harvested it after 4 hours, 1 day, 4 days, and 6 days. The use of the isotopic label allowed us to trace glucose-derived ^13^C into soil PLFAs and NLFAs. We expected the excess amount of labile carbon to trigger a strong response in microbial activity, where surplus carbon is distributed into storage compounds. Our hypothesis was that if bacterial NLFAs are necromass-markers derived from degraded phospholipids, they would show a much lower relative enrichment in ^13^C than their PLFA counterparts, as their majority would not originate from newly formed, ^13^C-labelled biomass. In contrast, if bacterial NLFAs are TAG-derived, bacterial NLFAs would show a similar or higher relative enrichment in ^13^C than their PLFA counterparts as glucose-¹³C would be allocated into newly formed storage compounds.

We found an immediate enrichment of ^13^C in fungi- and bacteria-specific PLFAs and NLFAs already four hours after glucose addition (Mann-Whitney test, P < 0.05; Fig. 6a). The ^13^C-enrichment increased for four days, and remained constant from day four to six. Fungal and gram-negative bacterial NLFAs showed higher relative ^13^C-enrichment compared to PLFAs throughout the experimental duration (t-test, P < 0.05). Conversely, gram-positive bacterial NLFAs were similarly enriched in ^13^C as PLFAs, except on day six. Fungal NLFAs showed consistently higher enrichment compared to bacterial NLFAs after the first day (t-test, P < 0.1). Most individual fatty acids showed higher ^13^C enrichment in NLFAs than PLFAs on day six (Fig. 6b). Of these, all fatty acids specific to general microbial biomass and fungi, two out of five gram-negative fatty acids (16:1ω7 and 17:0cy), and three out of six gram-positive fatty acids (16:0i, 17:0a, 17:0i) were more ^13^C-enriched in NLFAs than PLFAs (t-test, P < 0.05). None of the fatty acids were more ^13^C-enriched in PLFAs than NLFAs.

**Figure 6.**
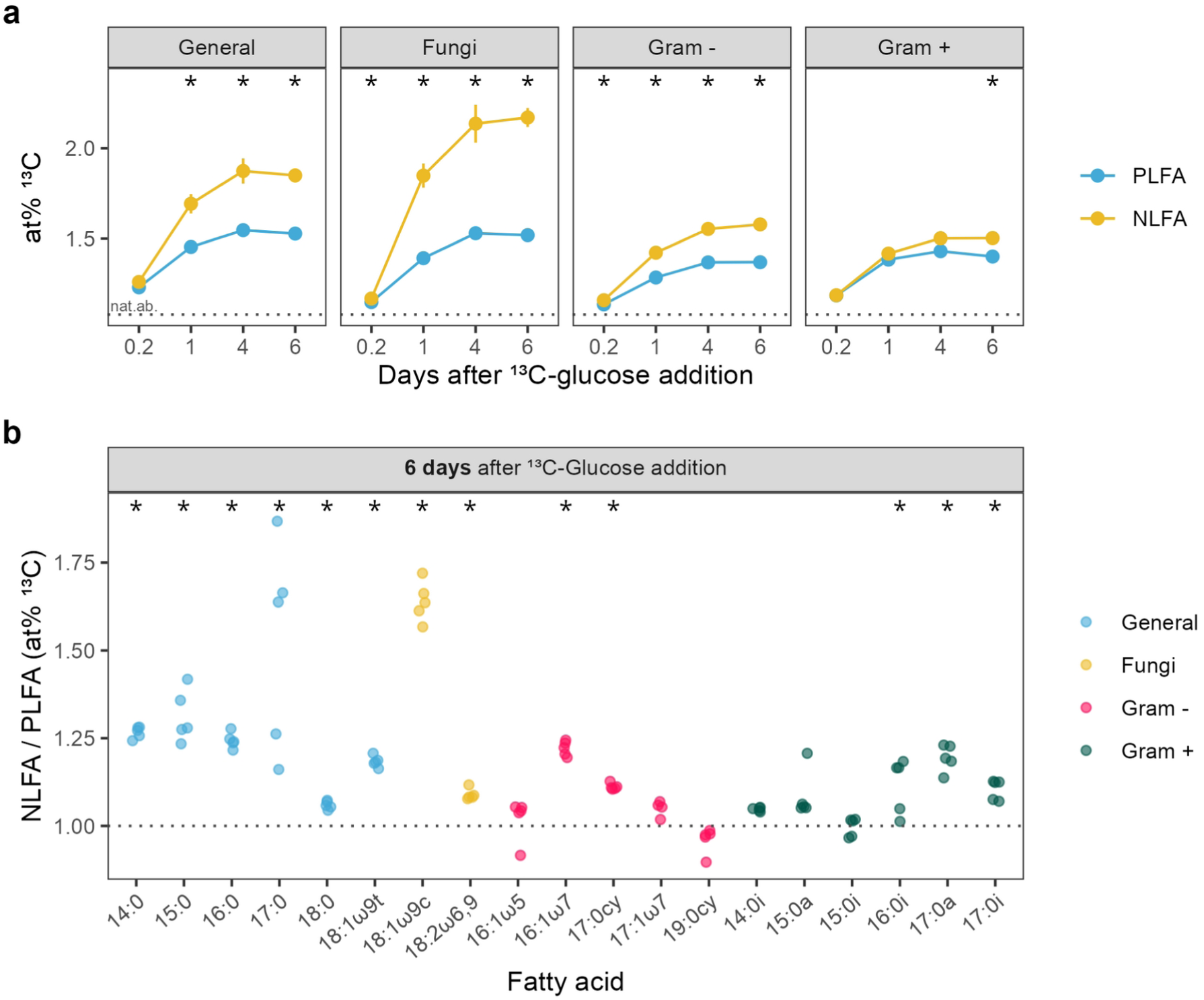
Incorporation of ^13^C into soil PLFAs and NLFAs after addition of ^13^C-glucose. **(a)** Time course of relative ^13^C incorporation into microbial PLFAs and NLFAs. Data is presented as the sum of all fatty acids specific to each respective microbial group. NLFAs tended to be more enriched than PLFAs. The dotted line represents average natural abundance ^13^C from control samples. Points represent means, and error bars represent the standard error. **(b)** NLFA/PLFA ratio of relative ^13^C incorporation into individual fatty acids, six days after the addition of ^13^C-glucose. Individual points represent replicate values. In both figures, asterisks indicate significant differences between PLFAs and NLFAs (t-test, p < 0.05). All fatty acids (PLFAs and NLFAs) in all time points were significantly enriched in ^13^C relative to unlabelled control values (Mann-Whitney test, p < 0.05). For a detailed description of the experimental setup and extraction procedure, see *Materials and Methods* section 8.3.

The rapid incorporation of added glucose-carbon into bacterial NLFAs, as well as the similar relative enrichment patterns of fungal and bacterial NLFAs, indicates bacterial accumulation of excess carbon in TAGs. This is similar to the findings of Rinnan et al. (2013) and Mason-Jones et al. (2023), who found allocation of labile-substrate-derived ^13^C into soil microbial NLFAs (without differentiating bacterial or fungal origin) within days after substrate addition. Previously we also observed that glucose addition to the same soil increased both fungal and bacterial NLFAs (Gorka et al., 2023b). Similarly, Koyama et al. (2018) found increased fungal and bacterial NLFAs, but not PLFAs, under elevated CO_2_ in plots with vegetation cover, suggesting that higher root exudation of labile carbon compounds increased microbial carbon storage. Thus, accumulating evidence from our research and others indicates that bacteria can accumulate TAGs in their biomass in response to certain environmental stimuli.

## 6. Contribution of intracellular DAGs to the NLFA pool

Intracellular DAGs, the anabolic precursors of both phospholipids and TAGs in cellular lipid synthesis, are a potentially third source of soil NLFAs. They can be formed through *de novo* synthesis via esterification of monoacylglycerol by acyltransferases, or turnover of TAGs, phospholipids, or phosphatidic acid (the intermediate compound between DAGs and phospholipids; Eichmann and Lass, 2015; Alvarez, 2016). The balance between these DAG formation pathways and further downstream DAG transformations (to membrane lipids or TAGs) determines the pool size of intracellular DAGs. Although DAGs are intermediate compounds in lipid synthesis, they can constitute a substantial fraction of intracellular neutral lipids: DAGs were found to occur in similar amounts (2–5 mol% of the total lipidome) as TAGs (2–8 mol%) in yeast (Ejsing et al., 2009; Klose et al., 2012). Since DAGs are also precursors for phospholipids, non-TAG-producing bacteria can still contribute to the soil neutral lipid pool. In these bacteria, fatty acid profiles from DAGs likely reflect PLFA profiles. Thus, even though data is lacking for bacteria, the potential presence of sizable intracellular DAG pools may explain the comparable amounts of detectable PLFAs and NLFAs in soil.

## 7. Conclusions and outlook

### 7.1 Bacterial NLFAs remain enigmatic—but data point towards storage compounds

We found several indications that bacterial NLFAs are predominantly derived from storage compounds: (i) the literature provides several observations of bacterial TAGs, (ii) a wide range of soil Actinobacteria and Proteobacteria are able to produce WS/DGAT enzymes catalysing the last step in TAG synthesis, and (iii) excess labile carbon is rapidly allocated into both soil bacterial NLFAs (similar to fungal NLFAs), suggesting usage of TAGs for surplus storage.

Nevertheless, we agree with the notion that bacterial NLFAs may be partly derived from sources other than storage compounds. Turnover of phospholipids from dead cells can result in neutral DAGs, which, as we show, is also recovered in the NLFA fraction (Table 2). A potential third source of NLFAs are intracellular DAGs, which are ubiquitous precursor compounds in phospholipid synthesis.

### 7.2 Outlook

Understanding whether NLFAs originate from carbon storage or necromass compounds requires identifying their source, either from TAGs or DAGs. In this regard, a disadvantage of the FAME extraction method is that the original lipid molecule containing the measured fatty acid can only be classified by its polarity, as structural information is lost once the fatty acid is cleaved. Shotgun lipidomics of bacterial pure cultures, where mass spectra of intact lipids are measured without prior chromatographic separation, can overcome this limitation (Ejsing et al., 2009; Klose et al., 2012; Wang et al., 2016). Furthermore, liquid chromatography (LC)-based approaches like ultra-high-pressure LC coupled with high-resolution mass spectrometry (UHPLC-HRMS) not only allow identification of intact lipid species (Bale et al., 2021; Couvillion et al., 2023)—and thus fingerprint TAGs and DAGs—but potentially enable absolute quantifications. Such lipidomics approaches could give valuable insight in the distribution of TAGs and DAGs not only in bacterial pure cultures, but also soil environments.

Experiments utilising stable isotopes could also elucidate the origin of bacterial NLFAs. For instance, addition of ^13^C-labelled TAGs and/or phospholipids and subsequent measurement of ^13^C incorporation in PLFAs, NLFAs, and respired CO_2_ could serve as a more comprehensive approach on the dynamics of retention, turnover, and microbial incorporation of lipids in soils. Experiments utilising stable isotopes which do not interfere with microbial metabolism (^2^H or ^18^O), could also give valuable insights into the origin of bacterial NLFAs. Canarini et al. (2023) reported that ^2^H in water vapour, which was in equilibrium with soil water, was detected in microbial PLFAs and NLFAs within 48 hours. Under a similar setup with higher temporal resolutions, simultaneous ^2^H enrichment in PLFAs and NLFAs would suggest NLFAs as storage markers, whereas a time-lag in ^2^H enrichment would indicate NLFAs to be derived from necromass as community turnover is required for the formation of phospholipid-derived NLFAs.

Full NLFA profiles are sensitive markers for soil carbon dynamics, but their enigmatic origin makes it difficult to define their functional role. Both suggested origins—storage compounds and necromass— potentially contribute to the final NLFA pool, as both TAGs and phospholipid-derived DAGs can elute in the neutral fraction in the FAME method. To date, soil studies have used bacterial NLFA mainly as necromass biomarkers. However, accumulating evidence from our research and others favours bacterial NLFA to be derived predominantly from storage compounds (i.e., TAGs). We suggest further experimental avenues to potentially quantify the contribution of each pathway to NLFA formation, such as coupling NLFA with stable isotopes or lipidomics approaches. Given the important role that NLFAs can have for soil biogeochemistry and microbial ecology, we hope that this perspective can foster new experimental avenues to quantify their source contribution as well as help researchers in guiding their interpretation of the function role of NLFAs in soil.

## 8. Materials and methods

### 8.1 Full NLFA profile literature search

A literature search was conducted on Web of Science using the search string *(NLFA* AND PLFA*) OR (“phospholipid fatty acid*” AND “neutral lipid fatty acid*”) OR (“lipid biomarker*” AND soil AND microbial)*. This resulted in a total list of 248 publications. Our selection criteria entailed that the publication has to show full PLFA and NLFA profiles (or grouped abundance data), and analyse soils. This yielded a filtered list of 15 publications. Plot data was extracted using PlotDigitizer (https://plotdigitizer.com/app). If the study used experimental treatments, only control values were extracted.

### 8.2 Extractions of soil and microbial pure culture PLFAs and NLFAs

Three soil types were harvested for the soil lipid extractions: a beech forest soil from Klausen-Leopoldsdorf, and a spruce forest as well as a grassland soil from Mariensee. All sites were located in Lower Austria, Austria. After removing aboveground biomass with a knife, the upper 5 cm of the soil were collected with soil cores (10 cm diameter) in replicates of five (n=5). Soils were homogenised by sieving (mesh size = 2 mm), and subsequently frozen, lyophilised, and stored at -20 °C until extraction.

Microbial pure culture strains were extracted as described in Gorka et al. (2023a; see also for complete descriptions of the microbial strains). These were gram-positive bacterial strains of Actinobacteria (*Solirubrobacter soli* DSM 22325, *Streptosporangium roseum* DSM 43021) and Firmicutes (*Paenibacillus alginolyticus* DSM 5050), gram-negative bacterial strains of Proteobacteria (*Labilithrix luteola* DSM 27648, *Chelativorans multitrophicus* DSM 6780, *Paraburkholderia xenovorans* DSM 17367), and fungal strains of arbuscular-mycorrhizal Glomeromycota (*Rhizophagus irregularis*), Ascomycetes (*Periconia macrospinosa* DSM 62880, *Trichoderma virens* DSM 3500), and Basidiomycetes (*Armillaria gallica* DSM 3732, *Psilocybe cyanescens* DSM 4999, *Lactarius subdulcis*, *Lactarius quietus*). All microbial pure cultures were lyophilised, and stored at -20 °C until extraction.

Total lipids were obtained with an adjusted FAME extraction method (Bligh and Dyer, 1959; Frostegård et al., 1991; Buyer and Sasser, 2012). In short, lyophylised samples were extracted with a monophasic solution of chloroform, methanol and citrate buffer (v:v:v = 1:2:0.8). A two-phase solution was reached by changing the solvent volume ratios via addition of chloroform and citrate buffer, allowing the collection of the lower phase containing the total lipids. These were subsequently fractionated via high-throughput silicic acid chromatography (Buyer and Sasser, 2012) with solvent of increasing polarity, yielding neutral lipids (chloroform and ethanol, v:v = 98:2; Drijber and Jeske, 2019), glycolipids (acetone), and phospholipids (chloroform, methanol and water, v:v:v = 5:5:1; Buyer and Sasser, 2012). Phospho- and neutral lipids were derivatised to FAME via mild alkaline methanolysis. The resulting PLFAs and NLFAs were injected on a gas chromatograph (7890B, Agilent Technologies) connected to a time of flight mass spectrometer (Tof-MS; Pegasus BT, LECO). Quantification was achieved via the internal standard methyl nonadecanoate. The exact specifications of the laboratory workflow are described in Gorka et al. (2023a).

Fatty acids below 0.5% of the total fatty acid abundance of each replicate sample were removed from further calculations. Taxonomic groups were calculated via the sum of the following fatty acids: Fatty acids were assigned as follows: General microbial biomass (14:0, 15:0, 16:0, 17:0, 18:0, 18:2ω9t), fungi (18:1ω9c, 18:2ω6,9, 18:3ω6,9,12), fatty acids with 20 carbon atoms (20:0, 20:1ω9, 20:2ω6,9, 20:3ω6,9,12, 20:4ω6,9,12,15, 20:5ω3,6,9,12,15), gram-negative bacteria (14:0-2OH, 14:1ω5, 16:0-2OH, 16:1ω7, 16:1ω9, 17:0cy, 17:1ω7, 17:1ω8, 18:1ω5, 19:0cy, 19:1ω9), gram-positive bacteria (14:0i, 15:0a, 15:0i, 16:0i, 17:0a, 17:0i, 18:0i), Actinobacteria (10Me-17:0, 10Me-18:0), and unspecified microbial biomass (15:1, 16:0-Me, 16:2ω6,9, 17:0-me). The fatty acid 16:1ω5, which is specific to both arbuscular-mycorrhizal fungi and gram-negative bacteria depending on the fatty acid type and ecosystem, was kept separate (Lekberg et al., 2022; Olsson and Lekberg, 2022).

### 8.3 ^13^C-labelled glucose addition experiment

The experimental setup was identical to Gorka et al. (2023b). In short, sieved (2 mm mesh size) beech forest soil (top 5 cm) was transferred to glass jars with a lid which allowed gas exchange. After one week acclimation time at 20 °C (the temperature was kept constant throughout the experimental duration), ^13^C-labelled glucose (99 at% ^13^C) was added (4 mg glucose g^−1^ soil) and homogenised via rigorous shaking, while control soils did not receive any treatment. Soil was subsequently harvested 4 hours, 1 day, 4 days, and 6 days after glucose addition (n=5). PLFAs and NLFAs were extracted as described in section 8.2; only for lipid fractionation, the pure solvents chloroform (neutral lipids), acetone (discarded), and methanol (phospholipids) were used. PLFAs and NLFAs were quantified on a gas chromatograph (Trace GC Ultra, Thermo Scientific) coupled to a mass spectrometer (ISQ, Thermo Scientific), and their isotopic ^13^C/^12^C ratios were measured on a gas chromatograph (Trace GC-Ultra, Thermo Fisher Scientific) coupled to an isotope ratio mass spectrometer (Finnigan Delta-V Advantage, Thermo Fisher Scientific) via a GC IsoLink (Thermo Fisher Scientific). For a complete description of the experimental setup see Gorka et al. (2023b). Fatty acids were assigned as follows: general microbial biomass (14:0, 15:0, 16:0, 17:0, 18:0, 18:1ω9t), fungi (18:1ω9c, 18:2ω6,9), gram-negative bacteria (16:1ω5, 16:1ω7, 17:0cy, 17:1ω7, 19:0cy), and gram-positive bacteria (15:0a, 15:0i, 16:0i, 17:0a, 17:0i).

### 8.4 Recovery of a pure DAG and wax ester

We used a diacylglycerol (1,2-dioleoyl-sn-glycerol; Avanti Polar Lipids) and wax ester (oleyl oleate; Sigma-Aldrich) with known attached fatty acids to test their recovery after lipid fractionation. Both compounds were subjected to lipid fractionation and derivatisation (mild alkaline methanolysis), and subsequently measured on a GC-Tof-MS system (as described in section 8.2). Recovery was calculated relative to replicates of the same compounds which did not undergo lipid fractionation.

### 8.5 DAGAT and WS/DGAT gene search

We searched the Joint Genome Institute’s (JGI) *Integrated Microbial Genomes with Microbiome Samples Expert Review* (IMG/M ER, tool ‘Function Search’; https://img.jgi.doe.gov/) for WS/DGAT (‘WS/DGAT C-terminal domain’, pfam06974), diacylglycerol acyltransferase (‘DAGAT’, pfam03982), and additionally for the PHA synthesis regulator PhaR (‘PHB/PHA accumulation regulator DNA-binding domain’, pfam07879; Maehara et al., 2002). The search was restricted to all JGI-sequenced bacterial isolates that occur in the specified ecosystem type ‘soil’. Species were classified based on their NCBI taxon identification numbers. The resulting dataset consisted of 2926 unique entries, of which 918 showed WS/DGAT, 74 DAGAT, and 790 PhaR. Phyla with very low species coverage (n < 20) were collectively denoted as ‘Other’ (i.e., Aquificae, Armatimonadetes, Candidatus Microgenomates, Cyanobacteria, Chloroflexi, Deinococcus-Thermus, Fusobacteria, Nitrospirae, Planctomycetota, Spirochaetes, Synergistetes, Verrucomicrobia).

### 8.6 Data analysis

All data were analysed with R (v4.3.1), with preliminary data transformations and calculations in LibreOffice Calc (v7.6.4.1). Statistical significance between groups were tested with the t-test, or if its assumptions were violated (Shapiro-Wilk’s test for normality and Levene’s test for homogeneity of variance), with the Mann-Whitney test. Plots were created with the package *ggplot2* (Wickham, 2016).

The cladogram was constructed using hierarchical taxonomic categories (domain, phylum, class, order, family, genus, species) via the function *as.phylo* from the package *ape* (v5.7-1; Paradis et al., 2004), and plotted with the package *ggtree* (v3.10.0; Wang et al., 2020b; Xu et al., 2021; Xu et al., 2022).

## Data availability

Open access to the dataset and R scripts to reproduce analyses and figures in the main article will be made available upon acceptance for publication on a public repository.

