## Supplementary Material for "Soil bacterial neutral lipid fatty acids: Markers for carbon storage or necromass?"

### Supplementary Figure and Tables

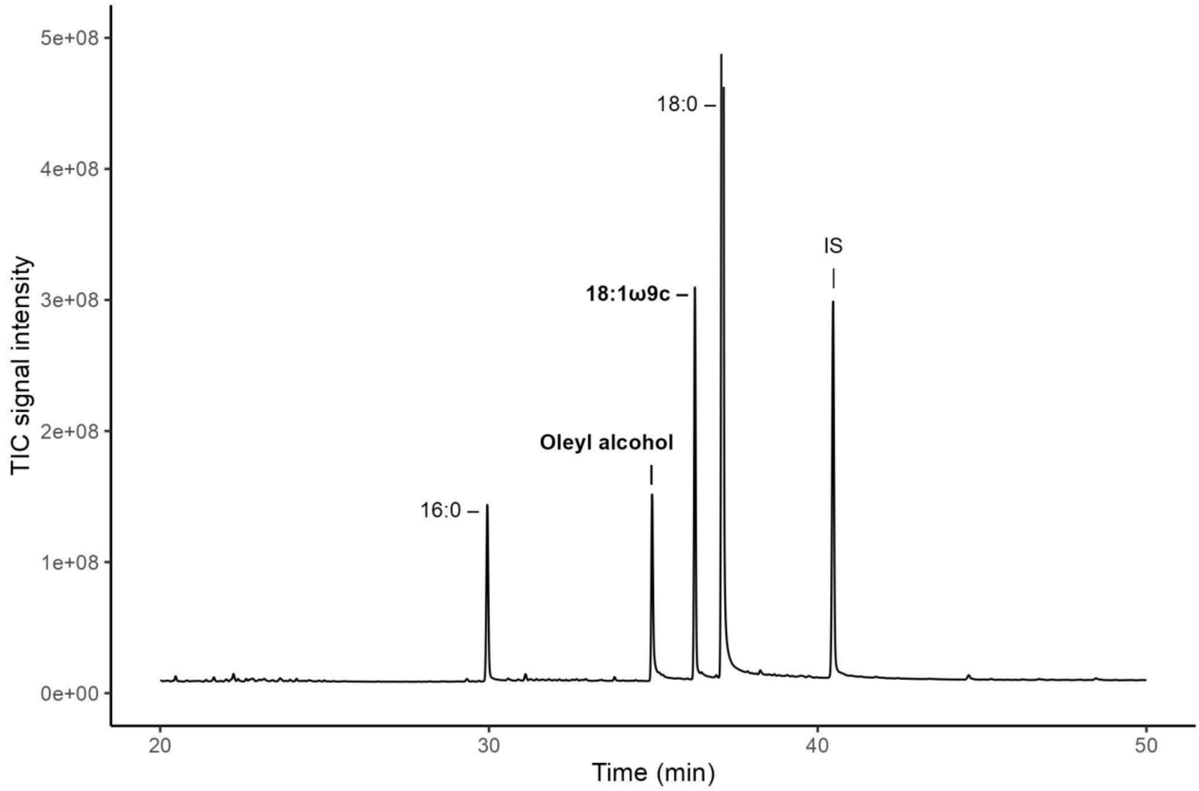


**Figure S1** Chromatogram of a methylated pure wax ester (oleyl oleate; NLFA fraction). Both moieties of the wax ester, i.e. the alcohol (oleyl alcohol) and methylated fatty acid (18:1ω9c) are measured in the same GC chromatogram with a standard temperature program for FAME measurements (Gorka et al., 2023a). Their recovery values are listed in Table 2. The fatty acids 16:0 and 18:0 are common contaminants in the neutral lipid fraction stemming from the plastic walls of silicic acid columns. IS, internal standard (FAME 19:0).

**Table S1** Individual PLFA and NLFA concentrations in nmol C $g^{-1}$ dm, and their respective NLFA/PLFA ratios, of the three analysed soils (see Fig. 1). NLFA/PLFA ratios above or equal 1 are marked in bold.

| **Specificity** | **FA** | **Beech forest** | | | **Pine forest** | | | **Grassland** | | |
|---|---|---|---|---|---|---|---|---|---|---|
| | | **PLFA** | **NLFA** | **NLFA/PLFA** | **PLFA** | **NLFA** | **NLFA/PLFA** | **PLFA** | **NLFA** | **NLFA/PLFA** |
| General | 14:0 | 18.5 | 76.8 | **4.1** | 39.9 | 176.8 | **4.4** | 35.1 | 91.8 | **2.6** |
| | 15:0 | 17.7 | 36.9 | **2.1** | 29.9 | 46.2 | **1.5** | - | 46.2 | - |
| | 16:0 | 355.2 | 873.8 | **2.5** | 691.7 | 933.1 | **1.3** | 690.4 | 1718.2 | **2.5** |
| | 17:0 | - | 23.1 | - | - | - | - | - | - | - |
| | 18:0 | 136.6 | 770.7 | **5.6** | 224.7 | 1734.1 | **7.7** | 228.5 | 999.2 | **4.4** |
| | 18:1ω9trans | 485.2 | 167.7 | 0.3 | 869.1 | 253.8 | 0.3 | 998.8 | 530.5 | 0.5 |
| Fungi | 18:1ω9cis | 322.9 | 852.6 | **2.6** | 843.4 | 901.1 | **1.1** | 692.2 | 1913.1 | **2.8** |
| | 18:2ω6,9 | 164.0 | 324.6 | **2.0** | 514.8 | 591.4 | **1.1** | 266.8 | 606.7 | **2.3** |
| | 18:3ω6,9,12 | - | - | - | - | - | - | - | 107.1 | - |
| 16:1ω5 | 16:1ω5 | 83.3 | 87.4 | **1.0** | 268.4 | 130.3 | 0.5 | 369.7 | 737.1 | **2.0** |
| C20 | 20:0 | - | 153.1 | - | - | 509.1 | - | - | 152.9 | - |
| | 20:4ω6,9,12,15cis | 20.6 | 17.7 | 0.9 | 45.3 | 39.2 | 0.9 | 47.0 | 62.8 | **1.3** |
| | 20:5ω3,6,9,12,15 | 18.1 | - | - | - | - | - | 40.2 | 53.0 | **1.3** |

**Table S1** (*continued*)

| Specificity | FA | Beech forest | | | Pine forest | | | Grassland | | |
|---|---|---|---|---|---|---|---|---|---|---|
| | | **PLFA** | **NLFA** | **NLFA/PLFA** | **PLFA** | **NLFA** | **NLFA/PLFA** | **PLFA** | **NLFA** | **NLFA/PLFA** |
| Gram - | 16:1ω7 | 200.6 | 172.3 | 0.9 | 468.9 | 178.5 | 0.4 | 534.7 | 1129.2 | **2.1** |
| | 16:1ω9 | 21.1 | - | - | 60.0 | - | - | - | - | - |
| | 17:0cy | 76.5 | 39.1 | 0.5 | 135.1 | 65.7 | 0.5 | 194.4 | 35.8 | 0.2 |
| | 17:1ω8 | - | 26.0 | - | - | - | - | - | 68.4 | - |
| | 18:1ω5 | 22.9 | - | - | 111.8 | - | - | 100.4 | - | - |
| | 19:0cy(ω9) | - | 33.5 | - | - | - | - | - | - | - |
| | 19:1ω9 | 433.7 | 52.0 | 0.1 | 505.1 | 131.2 | 0.3 | 507.5 | 54.3 | 0.1 |
| Gram + | 14:0i | - | 24.6 | - | - | - | - | 46.9 | - | - |
| | 15:0a | 117.2 | 68.1 | 0.6 | 188.9 | 96.0 | 0.5 | 340.3 | 140.4 | 0.4 |
| | 15:0i | 217.8 | 171.2 | 0.8 | 358.3 | 232.2 | 0.6 | 400.1 | 294.8 | 0.7 |
| | 16:0i | 75.2 | 162.3 | **2.2** | 110.3 | 181.9 | **1.6** | 107.7 | 142.8 | **1.3** |
| | 17:0a | 31.2 | 41.7 | **1.3** | 76.6 | 93.1 | **1.2** | 82.5 | 58.3 | 0.7 |
| | 17:0i | 35.0 | 34.7 | **1.0** | 59.3 | 47.3 | 0.8 | 78.8 | 61.6 | 0.8 |

**Table S1** (*continued*)

| Specificity | FA | Beech forest | | | Pine forest | | | Grassland | | |
|---|---|---|---|---|---|---|---|---|---|---|
| | | PLFA | NLFA | NLFA/PLFA | PLFA | NLFA | NLFA/PLFA | PLFA | NLFA | NLFA/PLFA |
| Actino | 10Me17:0 | 16.6 | 31.2 | **1.9** | - | 50.9 | - | - | - | - |
| | 10Me18:0 | 32.1 | 56.0 | **1.7** | 98.1 | 71.3 | 0.7 | 88.5 | 80.4 | 0.9 |
| unspecified | 16:0_Me | 19.5 | - | - | - | - | - | 59.6 | - | - |
| | 17:0_Me | 115.5 | 103.3 | 0.9 | 185.6 | 182.0 | **1.0** | 257.3 | 101.6 | 0.4 |

**Table S2** Occurrence of diacylglycerol acyltransferase (DAGAT) and wax ester synthase/diacylglycerol O-acyltransferase (WS/DGAT) specific genes in bacterial classes of soil isolate species. Both enzymes catalyse the last step in TAG synthesis. Asterisks indicate phylum assignment to 'Other' in the main figure and table.

| Phylum | Class | n | DAGAT (%) | WS/DGAT (%) |
|---|---|---|---|---|
| Acidobacteria | Acidobacteriia | 28 | - | 7.1 |
| | Blastocatellia | 1 | - | - |
| | Holophagae | 1 | - | - |
| | Vicinamibacteria | 1 | - | - |
| Actinobacteria | Actinomycetia | 1065 | 1.2 | 66.1 |
| | Rubrobacteria | 1 | - | 100 |
| | Thermoleophilia | 7 | - | 100 |
| Aquificae* | Aquificae | 2 | - | - |
| Armatimonadetes | Chthonomonadetes | 1 | - | - |
| Bacteroidota | Bacteroidia | 5 | - | - |
| | Chitinophagia | 49 | - | - |
| | Cytophagia | 81 | - | - |
| | Flavobacteriia | 78 | - | - |
| | Saprospiria | 1 | - | - |
| | Sphingobacteriia | 59 | - | - |
| Candidatus Microgenomates* | unclassified | 1 | - | - |
| Chloroflexi* | Ktedonobacteria | 3 | - | - |
| | unclassified | 2 | - | - |
| Cyanobacteria* | unclassified | 15 | 86.7 | - |
| Deinococcus-Thermus* | Deinococci | 15 | - | - |
| Firmicutes | Bacilli | 386 | 0.3 | 0.3 |
| | Clostridia | 43 | - | 2.3 |
| | Negativicutes | 2 | - | - |
| | Tissierellia | 2 | - | - |
| | unclassified | 1 | - | - |

**Table S2** *(continued)*

| Phylum | Class | n | DAGAT (%) | WS/DGAT (%) |
|---|---|---|---|---|
| Fusobacteria* | Fusobacteriia | 1 | - | - |
| Nitrospirae* | Thermodesulfovibrionia | 1 | - | - |
| Planctomycetota* | Planctomycetia | 4 | - | - |
| Proteobacteria | Acidithiobacillia | 3 | - | - |
| | Alphaproteobacteria | 335 | - | 9.9 |
| | Betaproteobacteria | 287 | - | 17.1 |
| | Deltaproteobacteria | 90 | 48.9 | 73.3 |
| | Epsilonproteobacteria | 3 | - | - |
| | Gammaproteobacteria | 333 | 0.9 | 16.2 |
| | Oligoflexia | 3 | - | - |
| Spirochaetes* | Spirochaetia | 7 | - | - |
| Synergistetes* | Synergistia | 1 | - | - |
| Verrucomicrobia* | Methylacidiphilae | 1 | - | - |
| | Opitutae | 1 | - | - |
| | Spartobacteria | 1 | - | - |
| | Verrucomicrobiae | 5 | - | - |
